# Bet-hedging across generations can affect the evolution of variance-sensitive strategies within generations

**DOI:** 10.1101/600452

**Authors:** Thomas R. Haaland, Jonathan Wright, Irja I. Ratikainen

## Abstract

In order to understand how organisms cope with ongoing changes in environmental variability it is important to consider all types of adaptations to environmental uncertainty on different time-scales. Conservative bet-hedging represents a long-term genotype-level strategy that maximizes lineage geometric mean fitness in stochastic environments by decreasing individual fitness variance, despite also lowering arithmetic mean fitness. Meanwhile, variance-prone (aka risk-prone) strategies produce greater variance in short-term payoffs because this increases expected arithmetic mean fitness if the relationship between payoffs and fitness is accelerating. Using two evolutionary simulation models, we investigate whether selection for such variance-prone strategies are counteracted by selection for bet-hedging that works to adaptively reduce fitness variance. We predict that variance-prone strategies will be favored in scenarios with more decision events per lifetime and when fitness accumulates additively rather than multiplicatively. In our model variance-proneness evolved in fine-grained environments (with lower correlations among individuals in energetic state and/or in payoffs when choosing the variable decision), and with larger numbers of independent decision events over which resources accumulate prior to selection. In contrast, geometric fitness accumulation caused by coarser environmental grain and fewer decision events prior to selection favors conservative bet-hedging via greater variance-aversion. We discuss examples of variance-sensitive strategies in optimal foraging, migration, life histories and cooperative breeding in light of these results concerning bet-hedging. By linking disparate fields of research studying adaptations to variable environments we should be more able to understand the effects in nature of human-induced rapid environmental change.

**Data deposition:** R code is available upon request.

## Introduction

The world is a stochastic place and evolution favors organisms that are able to persist in the face of such random variation within and among lifetimes in resource availability, predation risk or environmental conditions (Cohen 1966; Simons 2002; Botero et al. 2015). While various adaptations to stochasticity have been the topic of intense research interest in many different scientific fields, including behavioral ecology, physiology and evolutionary ecology, much of this research unfortunately remains rather disparate with little unifying work being done (but see Haaland et al. 2019).

Ever since the work of Thomas Caraco and colleagues in the 1980s, it has been widely accepted in behavioral ecology that animals should exhibit variance-sensitive behavior (Caraco et al. 1980; Caraco 1981; Real and Caraco 1986), aka. ‘risk-sensitivity’ (Smallwood 1996; Stephens et al. 2007). Arriving at a time when optimality models were gaining traction in evolutionary and behavioral ecology, this important theoretical development highlighted the crucial point that the optimal strategy can also depend upon the variation in payoffs around the mean. This is because the payoffs from a specific behavior (e.g. obtaining successive food items) do not necessarily relate linearly to fitness - i.e. utility functions are in most cases expected to be non-linear (Caraco et al. 1980; Stephens 1981). For example, the fitness benefits of a food resource are expected to increase exponentially early on when there is a real danger of starvation, and they will flatten out towards an asymptote due to diminishing returns of additional resource gains when the animal becomes satiated (see Fig. 1 in Wright and Radford 2010). This is a general property of some threshold level of resources being needed to complete a certain task, be it surviving a cold night, undergoing migration, achieving a certain social dominance rank, attracting a mate or successfully raising offspring (Real and Caraco 1986; Bednekoff 1996; Kacelnik and Bateson 1997; Hurly 2003; Ratikainen et al. 2010). Importantly, when the relationship between resources gained and fitness is accelerating (i.e. the utility function is convex), a variance-prone strategy resulting in variable resource gain provides higher expected fitness. Conversely, if the organism’s utility function is concave (decelerating), being variance-averse and preferring less variable sources of resource gain provides higher expected fitness (see Fig. 1 in Wright and Radford 2010). These predictions from the energy budget rule (Caraco 1981; Stephens 1981; Bednekoff 1996) have largely been supported in experimental studies of foraging decisions in a range of animal species (Shafir 2000) and recently even plants (Dener et al. 2016), but results are inconsistent, indicating that there are still unresolved issues in this paradigm (Kacelnik and El Mouden 2013).

**Figure 1:**
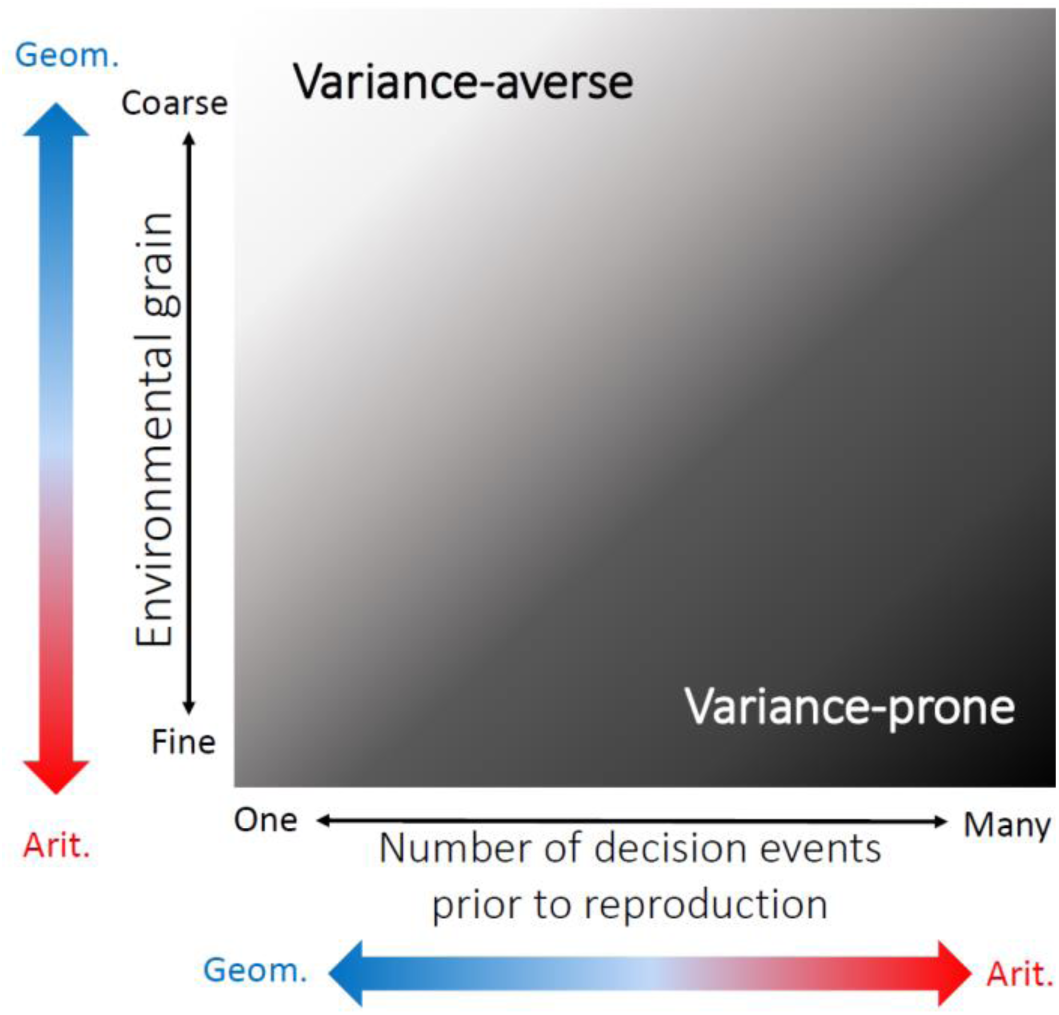
Conceptual illustration of the factors predicted to affect selection for variance-averse (light) versus variance-prone (dark) strategies in light of bet-hedging theory. An increasing number of decision events prior to reproduction (moving right on the x-axis) across which payoffs accumulate additively, and a finer environmental grain (moving down on the y-axis) which decreases correlations in payoffs among variance-prone individuals, should in combination favor variance-prone strategies, since they both shift the balance from geometric to arithmetic fitness accumulation (colored arrows). In contrast, fewer decision events prior to reproduction and coarser environmental grain causes fitness to accumulate geometrically over time, favoring variance-averse strategies providing lower variation in fitness despite also providing lower expected fitness.

In a parallel field of study in evolutionary ecology, bet-hedging has been defined as a strategy increasing its probability of fixation in the population through decreasing the variation in fitness across generations despite also decreasing mean fitness (Slatkin 1974; Kaplan and Cooper 1984; Seger and Brockmann 1987; Starrfelt and Kokko 2012). The success of a bet-hedging strategy lies in reproduction across generations being an inherently multiplicative process, so the success of a lineage over time is best estimated by geometric mean fitness across generations rather than the arithmetic mean (Cohen 1966; Lewontin and Cohen 1969; Philippi and Seger 1989). Geometric means are much more sensitive to variation than are arithmetic means, and thus a genotype experiencing less variation in fitness across generations can spread despite having a lower expected fitness in any one generation (Simons 2011; Starrfelt and Kokko 2012). This concept was first used to explain seed dormancy of desert annuals (Cohen 1966; Clauss and Venable 2000; Gremer and Venable 2014), but has received an upsurge of attention in recent years as it provides a tantalizing explanation for a range of seemingly ‘suboptimal’ strategies observed in bacteria, animals, plants and fungi (Simons 2002; Simons and Johnston 2003; Balaban et al. 2004; Einum and Fleming 2004; Beaumont et al. 2009; Salathé et al. 2009; Childs et al. 2010; Graham et al. 2014; Botero et al. 2015; Haaland et al. 2019). Theory has shown that bet-hedging is most important when environmental fluctuations between generations are larger than those within generations, i.e. in ‘coarse-grained’ environments, and more generally when temporal environmental fluctuations become more important than spatial environmental fluctuations (Starrfelt and Kokko 2012; Crowley et al. 2016; Haaland et al. 2019).

Unfortunately, the term bet-hedging is often misused and misunderstood, leading to considerable confusion in the literature. For instance, the genotypic strategy of diversified bet-hedging (DBH) which produces phenotypically different individuals can be adaptive in an unpredictably fluctuating environment because it lowers the fitness correlations among individuals, leading to a lower variance in fitness at the genotype level (Bull 1987; Seger and Brockmann 1987; Starrfelt and Kokko 2012). However, DBH is often invoked to explain any observed phenotypic variation in a trait, without checking whether this reduces genotype-level fitness variance at the expense of arithmetic mean fitness (Olofsson et al. 2009; Rees et al. 2010). Alternatively, a genotype may lower its variation in fitness through ensuring that each individual experiences low variation in fitness, i.e. a conservative bet-hedging (CBH) strategy. Such a strategy is often manifest as ‘playing it safe’ in the face of uncertainty, e.g. due to predation or starvation risk (Simons 2011; Haaland et al. 2019), but in order for this to be termed bet-hedging a reduction in expected fitness at the individual level is also required. Otherwise playing it safe or ‘insurance’ is simply the optimal strategy, and bet-hedging is not required as an explanation. In this respect, variance-averse decisions are superficially similar to a CBH strategy (such that they are actually often confused in the literature), in that variability is adaptively avoided. This link was mentioned almost 30 years ago by Frank and Slatkin (1990), but they only considered the choice of variance-aversion by foragers in the concave (decelerating) part of the utility function. In such a situation, variance-aversion not only lowers an individual’s variance in fitness, but also increases its average fitness, i.e. the strategy does not increase geometric mean fitness at the cost of a lower arithmetic mean fitness, and does thus not constitute bet-hedging (Seger and Brockmann 1987; Starrfelt and Kokko 2012). However, once one considers bet-hedging in the long-term alongside any variance sensitive strategy, even in the convex (accelerating) part of the utility function, variance-aversion might still be favored as a CBH strategy. This is because although variance-proneness here would cause an increase in an individual’s average fitness, there may be bet-hedging benefits to being variance-averse, i.e. achieving lower fitness variation may be favored in the long term at the genotype level despite lowering average individual fitness in the short term.

While well-known in economics, biologists have typically only applied variance-sensitivity theory to foraging behavior. However, the choice between a safe option giving predictably moderate rewards and a variable option giving small or large rewards applies to any number of problems in behavioral ecology and other realms of biology, including group formation and optimal group sizes (Clark and Mangel 1986; Poethke and Liebig 2008), the evolution of cooperation (Rubenstein 2011; Kennedy et al. 2018) and reproductive decisions such as alternative mating strategies (Alonzo and Warner 2000; Carter et al. 2015), optimal litter sizes (Mountford 1968; Boyce and Perrins 1987), or biasing investment towards male versus female offspring in species with high reproductive skew (Clutton-Brock et al. 1981; Røed et al. 2007). In cases where the utility function is accelerating, choosing the variable option will provide considerably higher mean fitness, but it may also increase the variance in mean fitness, which would then have bet-hedging consequences. Importantly, any fitness variance created by such variance-prone decisions will decrease over the course of a greater number of events in a lifetime, if payoffs accumulate additively. Therefore, the potential for bet-hedging advantages from lowering fitness variance will be reduced the more variance-sensitive decisions are made prior to selection (Fig. 1). Furthermore, this reduction in fitness variance at the individual level will be more important in terms of genotype-level geometric mean fitness if the correlations in payoffs among variance-prone individuals are high (coarse-grained environment). As we show in this paper, the adaptive nature of short-term variance sensitivity (especially outside of foraging behavior) needs to be considered in light of long-term bet-hedging strategies and consequences.

Here we present two individual-based simulation models of individuals facing a decision between options providing constant versus variable rewards. We explore evolutionary outcomes in scenarios differing in their grain of environment and in the number of decision events made within a lifetime prior to selection. These different scenarios can be envisioned as modelling traits related to different activities. For example, simulations with many decision events prior to selection can represent scenarios for traits that contribute additively to reproductive success, such as foraging-related decisions where payoffs accumulate over a long sequence of behavioral events. However, simulations with a small number of decision events per lifetime would represent traits/decisions related more directly to reproductive events, such as clutch size or timing of breeding. Here, the payoffs for decisions do not simply accumulate additively across events within a lifetime, but will more obviously accumulate multiplicatively across consecutive generations in a lineage. We therefore predict the evolution of variance-averse behaviors despite their lower (arithmetic) mean fitness benefits in scenarios favoring bet-hedging strategies – i.e. coarse-grained environments and a low number of decisions prior to selection (Fig. 1). This analysis provides novel links between short-term behavioral and long-term evolutionary adaptations to environmental variability.

## Model description

We model populations of females that choose between a risky versus a safe strategy to obtain fitness rewards, which we refer to respectively as ‘variable’ versus ‘constant’ rewards. The ‘variable’ payoff is determined by some randomly varying environmental factor (‘good’ versus ‘bad’ conditions), whilst the ‘constant’ payoff is always the same. We hypothesize that the importance of arithmetic versus geometric mean fitness in determining long-term evolution of these two strategies should depend upon: (i) the number of decision events or times (termed *n*) that the trait is used to gain resources prior to selection; and (ii) the ‘grain’ of the environment, *g* - i.e. the extent to which the environmental fluctuations affect the individuals in the population in the same way (see Introduction and Fig.1). We use two different versions of our simulation model to investigate these effects.

### Model 1: Risky versus safe strategies when payoffs accumulate additively versus multiplicatively

In Model 1, the only effect of the environment is whether the individuals using the variable strategy get a high or low payoff. We simply set the conditions to be good (1) or bad (0) with equal probability, 0.5. Environmental grain *g*_*r*_(for ‘grain of resources’) is incorporated as the correlations in resources among patches, which in this model determines the payoffs of individuals choosing the variable strategy. If *g*_*r*_ = 1, the quality of the variable patch is the same for all individuals, whereas if it is *g*_*r*_*=* 0, each individual choosing the variable strategy has an independent chance of encountering a patch that is rich or low in resources, each with probability 0.5. For intermediate *g*_*r*_, the overall patch state *R* is first calculated, after which the probability of each individual *i* experiencing environment *r*_*i*_ is determined by P(*r*_*i*_ = *R*) = 0.5 + (*g*_*r*_/2) and P(*r*_*i*_≠ *R*) = 1 - P(*r*_*i*_ = *R*). We acknowledge that some combinations of environmental grain and number of decision events are more realistic than others. For example, the success of risky reproductive strategies is typically determined by weather conditions experienced by at least a considerable part of the population, making low *n* and high *g*_*r*_ an interesting scenario. For foraging-related traits such as choosing between variable versus constant patches, the environment represents patch quality, and is probably best approximated with a low *g*_*r*_ (individuals choosing variable patches are likely to differ in their success in any given time step) and high *n* (foraging happens many, many times per lifetime, for most species). Yet, we here examine the full range of combinations of *n* = {1, 2, 5, 10} and *g*_*r*_ = {0, 0.25, 0.5, 0.75, 1}.

In order to investigate the importance of bet-hedging, we set the payoff from the constant strategy to be *W*_*const*_ = *µ*(1-*a*), so that the proportion *a* ϵ [0,1) represents the penalty for choosing the constant strategy relative to the expected payoff of the variable strategy, which is *µ*. We set the variability in payoffs of the variable strategy to be the proportion *b* ϵ (0,1] of *µ*, such that the payoffs are *W*_*var*_(*r*_*i*_ = 0) = *µ*(1-*b*) or *W*_*var*_(*r*_*i*_ = 1) = *µ*(1+*b*). If *b* = 1, we note that bad outcomes provide payoff *µ*(1-1) = 0, whereas if *b* = 0 the payoffs are equal. Intuitively, a larger *b* provides a larger punishment in terms of geometric mean fitness and would thus require a larger *a* (the penalty in payoff of the constant strategy relative to the mean payoff for the variable strategy) for the variable option to still be favored. In this model, we expect the constant strategy to be favored when fitness accumulation is entirely multiplicative across generations, as is the case when *g*_*r*_ = 1, because the arithmetic mean fitness advantage of a lineage playing a risky strategy should be outweighed by the reduction in geometric mean fitness created if mean fitness of the genotype varies more across generations (top left in Fig. 1). This variability increases as *g*_*r*_ approaches 1, since all individuals experience the same environmental condition (high correlation in fitness among individuals), and decreases as the number of decision events or time steps *n* becomes larger, since the variance in lifetime average payoff (arithmetic mean of a series of Bernoulli decision events) decreases with more decision events. Specifically, the geometric mean fitness of a lineage playing the variable strategy with *n* decision events in each individual’s lifetime is given by:

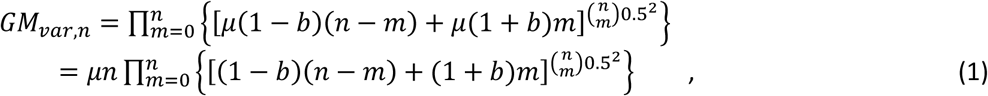

where *m* is the number of ‘successful’ decision events (i.e. for how many of the *n* times the environment was of good quality).

Comparing this to the payoff of playing the constant strategy *n* times, which is *µn*(1-*a*), we can identify a condition for a value of *a* below which the constant strategy should be favored (Figure 2): 

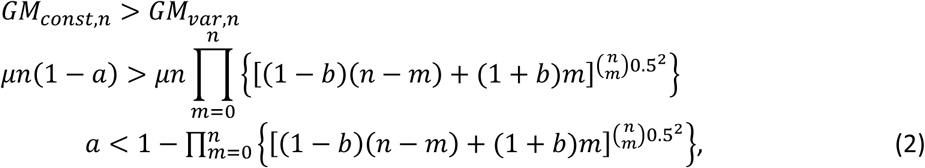

which does not depend on *µ*, but does depend on *n*, with this condition becoming less stringent as *n* increases. Figure 2 shows the results from inequality (2) for a range of values of *b* and *n*, such that the values represent the maximum value of *a* (reduction in mean payoffs from choosing the constant strategy) favoring the constant strategy. We use this to choose suitable values of *a* and *b* so that simulated populations are predicted to switch from safe to risky strategies as *n* increases and *g*_*r*_ decreases. For the simulation results shown here, we use *a* = 0.1, *b* = 0.9 and *µ* = 2 (i.e. the constant strategy provides payoff 0.9*2 = 1.8, and the variable patch provides payoff 2*0.1 = 0.2 or 2*1.9 = 3.8, each with probability 0.5). These parameter values give approximately equal fitness for the two strategies when *n* = 5 (follow contour line for *a* = 0.1 up to *b* = 0.9 in Fig. 2), thus favoring the constant strategy for lower values of *n* and the variable strategy for higher values of *n*. However, note that we predict this balance point to shift as *g*_*r*_ decreases from 1, since this entails a shift from multiplicative to additive long-term fitness accumulation (cf. Fig. 1).

**Figure 2:**
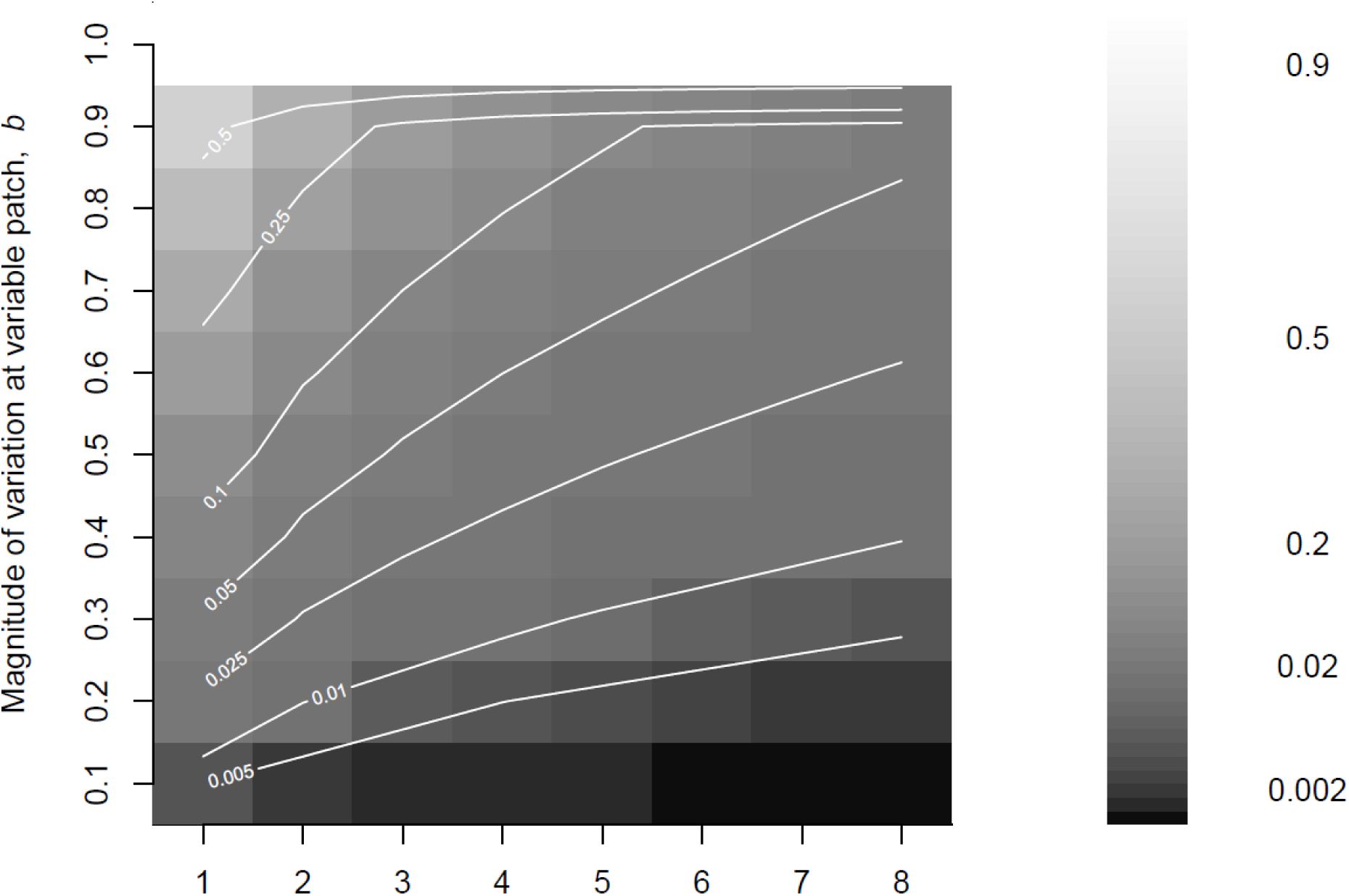
Maximum proportional decrease in the mean payoffs at the constant patch relative to the variable patch (*a*, shown in the shade of grey background and contour lines) that allow the constant patch to be favored – i.e. the upper bound of *a* for choosing the constant option. Values are shown for different amounts of variation at the variable patch, *b*, and number of decision events prior to reproduction, *n*. Limits of *a* in each grid cell are calculated by solving inequality (2) for the given value of *b* and *n*.

### Model 2: Risky versus safe strategies with differences in individual energetic state

Model 2 follows the same structure as Model 1 (above), but it explicitly also includes individual energetic state, *x*, and a sigmoid utility function relating energetic state to fitness,

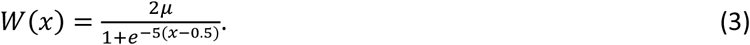

Individual state was an unnecessary complication for Model 1, but here it allows us to capture the mechanisms typically associated with the theory and empirical assessments of variance-sensitivity (see Introduction). We now allow the environment to vary continuously between 0 (bad) and 1 (good). The overall environment *E* determines the energetic state of individuals, which are stochastically drawn from a uniform distribution around *E*. The width of this distribution is given by the grain of the environment *g*_*e*_, with the lower boundary set to *E* - (1-*g*_*e*_)\**E* and the upper boundary to *E* + (1-*g*_*e*_)*(1-*E*). Thus, if *g*_*e*_ = 1, all individuals have the same energetic state of *x*_*i*_ = *E*, and if *g*_*e*_ = 0, individual states are drawn randomly from between 0 and 1. We consider the environment in such cases to represent something like overnight roost temperature determining mass loss for birds metabolizing fat to stay warm (Brodin 2007), which might vary considerably among individuals due to differences in nest insulation or location, and thus have a lower *g*_*e*_. Alternatively, an environmental variable such as winter harshness determining body condition at the beginning of the next season (Monteith et al. 2013) is likely to affect everyone equally and to have a higher *g*_*e*_.

Once individual states are determined, we allow variance-sensitive individuals that ended up in low state (*x* less than 0.5, which is the inflection point of *W*(*x*)) to forage at a variable patch, which will increase (if successful) or decrease (if unsuccessful) their energetic state by 0.1. The probability of success at the variable patch is determined by the overall state of the resources, *R*, and the grain of the resources, *g*_*r*_. These act in the same way as in Model 1. We assume that variance-averse individuals and individuals in high energetic state forage at a constant patch with moderate returns, which provides enough food to predictably keep them on the same energy level (i.e. leaving their state unchanged). Finally, updated energetic state *x’* is converted to resources used to acquire fitness, as determined by the utility function *W*(*x’*). Scaling the function to an upper asymptote of 2\**µ* allows a similar interpretation of *µ* as in Model 1, so that *µ*/*n* represents mean reproductive rate.

### Simulation algorithm

We use individual-based simulations to investigate the fate of a gene determining its bearer’s probability of playing a variance-sensitive versus an all variance-averse strategy. After *n* time steps or decision events in a lifetime where individual *i* gathers resources using a variable or constant strategy prior to reproduction (as described for the two model versions above), offspring are produced proportionally to the total amount of resources the individual gathered, *W*_*i*_ = Σ_*t*=1->*n*_W_*t,i*_. Then, depending upon the number of offspring produced and the population density (determined by between-year mortality *α* and adult population size *N* relative to the carrying capacity *K*), a number of recruits to next year’s population are chosen at random from the pool of offspring. This number is Σ_*i*_*W*_*i*_, if Σ_*i*_*W*_*i*_+ *N*α* > *K*, and *N*\**α* - *K* if Σ_*i*_*W*_*i*_+ *N*α* < *K*. Thus, we assume that juveniles are not affected by density regulation, that offspring of adults that die overwinter are able to survive without their parents, and that there may be a period over the course of the winter where the total population size exceeds *K*, but that by the time the next season begins *N* ≤ *K*. Since between-year mortality is random with respect to the effects of the gene of interest, this does not affect the evolutionary outcome. A proportion *m*_*p*_ of offspring produced are then subject to mutation, which changes their gene value according to a Gaussian distribution around the parent’s gene value, with standard deviation *m*_*σ*_. Finally, since the gene determines probabilities, gene values are constrained to be between 0 and 1, such that negative values are set to 0 and values exceeding 1 are set to 1.

Simulations were run for 2000 seasons and 100 independent replicates for each parameter combination were produced. Populations were initiated with uniformly distributed gene values between 0 and 1, and with an initial population size *N*_0_ at the carrying capacity *K*, which was set to 5000 individuals. For the results presented here, we use *m*_*p*_ = 0.005, *m*_*σ*_ = 0.05, and *µ* = 2, but varying these parameters did not affect our conclusions. The model is coded in R version 3.3.1 (R Core Team 2016) and the code is available upon request.

## Results

### Model 1

Figure 3 shows the evolved gene values at the end of the simulations for discrete (between-year mortality *α* = 1) and overlapping (*α* = 0.5) generations in scenarios with different grains of resources, *g*_*r*_, and number of decision events prior to reproduction, *n*. All populations survived until the end of the simulations and maintained stable population sizes at carrying capacity. Gene values stabilized and were highly repeatable across replicate simulations (as evidenced by the small error bars). Whether generations were overlapping or discrete did not affect evolutionary trajectories (as seen by the strong similarity among panels in Fig. 3). There is a strong interaction effect of *g*_*r*_ and *n* on the gene values for probability of playing the variable strategy. Specifically, when *g*_*r*_ is low (no/low correlations among the payoffs of individuals playing the variable strategy), the variable strategy is favored regardless of *n*. In contrast, for higher values of *g*_*r*_, low values of *n* strongly favor the safe strategy, whereas simulations with higher values of *n* tend less strongly towards the safe strategy as *g*_*r*_ increases. Indeed, for *n*=10, the populations ended up choosing the variable strategy nearly 100 % of the time no matter what the among-individual correlations in payoffs, and when *g*_*e*_ =0 or 0.25 this was the case for all values of *n*. Note that for *n*=1 (red line, open circles, Fig. 3), this result at low *g*_*r*_ involves half of the population hardly getting any reproductive success at all, but this does not prevent the spread of the gene for choosing the variable strategy, since the low correlations in fitness among individuals lead to low variance in fitness across generations at the genotype level.

**Figure 3:**
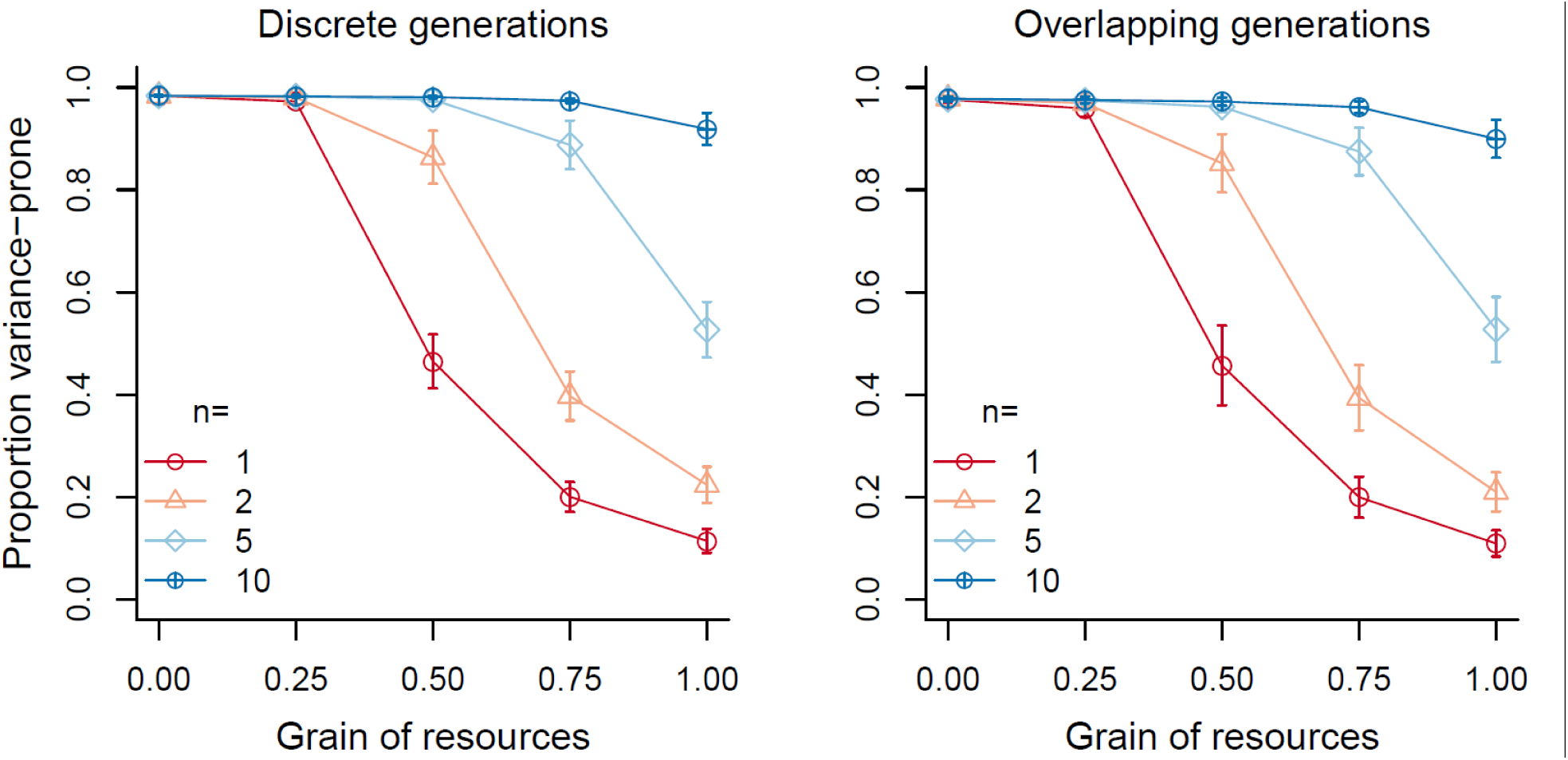
Mean evolved gene values for the proportion of the population playing variance-prone strategies at the end of the simulations for discrete (left panel, between-year mortality *α*=1) and overlapping (right, *α*=0.5) generations, for different resource grain (*g*_*r*_, x-axis) and number of decision events prior to reproduction (colors, point types). Points indicate means and error bars indicate standard deviations across 100 replicate populations, and relative point size represents the proportion of populations surviving until the end of the simulation (for the parameters shown here, all populations survived). Payoffs of the variable and constant strategies are determined as described in Model Description section for Model 1, using the parameter values *µ*=2, *a*=0.1 and *b*=0.9.

### Model 2

Figure 4 shows the evolved gene values at the end of the simulations for overlapping and discrete generations in scenarios with different grains of environment (*g*_*e*_), grains of resources (*g*_*r*_), and number of decision events prior to reproduction (*n*). In the Model 2 simulations, population size was generally more variable and extinction rates were higher than in Model 1 (above), especially in the simulations with low *n*, discrete generations and a large grain of environment *g*_*e*_ (i.e. strong correlations among individual energetic states). Grain of resources *g*_*r*_ (the correlation among payoffs of individuals choosing the variable strategy) did not affect extinction risk – i.e. extinction risk increases from left to right in each subpanel, but not across subpanels in Fig. 4. We see high frequencies of variance-sensitive strategies when either *g*_*e*_ or *g*_*r*_ were low, and selection for variance-sensitivity is consistently stronger for low *n*. This is shown by the red and orange lines being significantly higher than blue lines in the leftmost subpanels of the top row of Fig. 4. In addition, the red line is significantly higher than other lines in the leftmost subpanels in the bottom row of Fig. 4. Variance-aversion is increasingly favored as *g*_*r*_ approaches 1, but only for *n*=1 (and to a limited extent when *n*=2) and when there is a high *g*_*e*_. This effect appears clearest in populations with overlapping generations, but this is due to the discrete generation populations almost all going extinct when *g*_*e*_ =1 and *n* = 1. Evolution of the gene for variance-sensitivity is completely unaffected by *g*_*e*_ or *g*_*r*_ when *n* is sufficiently high (the slope of the blue lines is zero and their elevation is unchanged across panels in each row of Fig.4).

**Figure 4:**
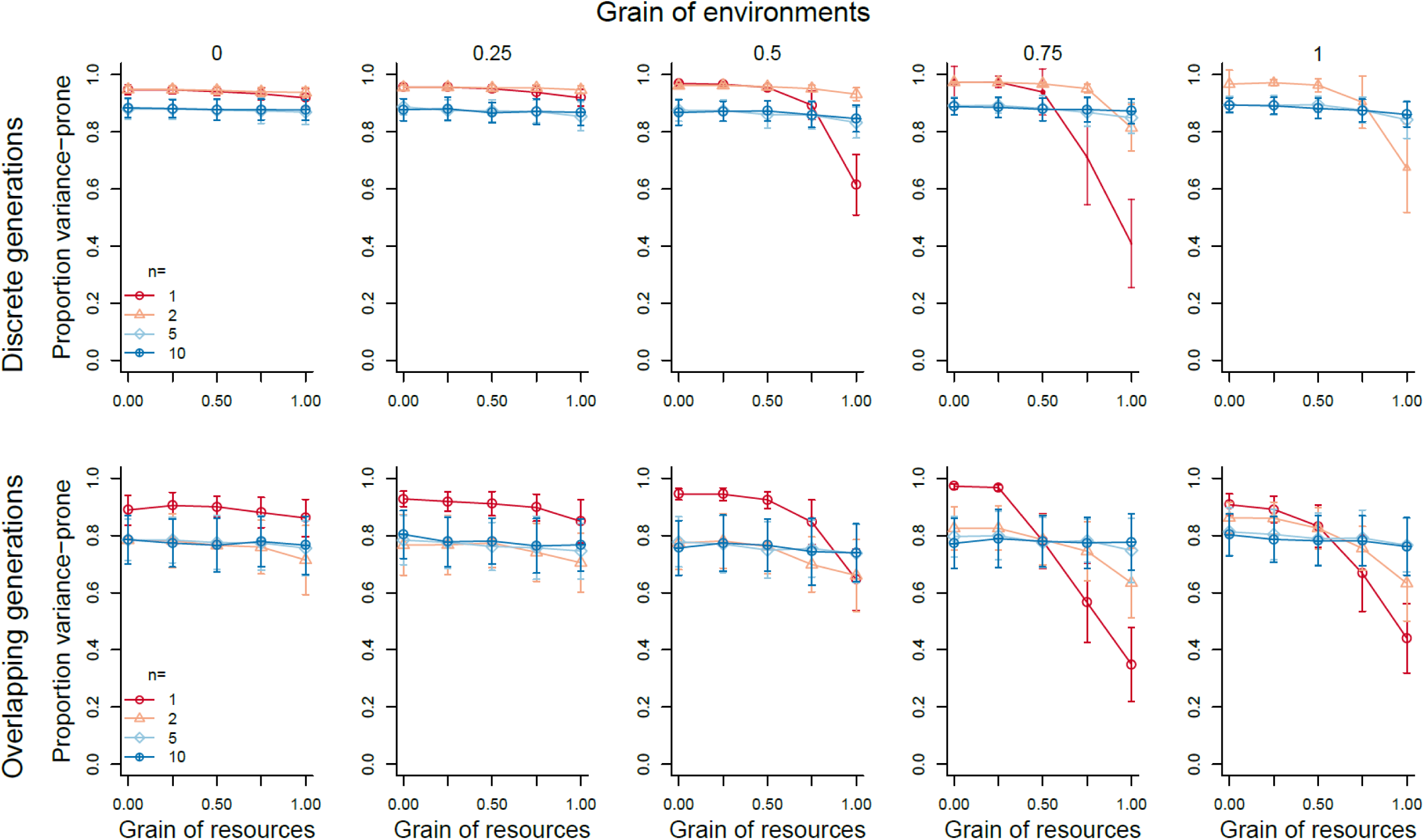
Mean evolved gene values for the proportion of the population playing variance-prone strategies at the end of the simulations for discrete (top row, between-year mortality *α*=1) and overlapping (right, *α*=0.5) generations, for different grain of environment (*g*_*e*_, x-axes), grain of resources (*g*_*r*_, given by the number 0 to 1 above each column) and number of decision events prior to reproduction (colors, point types). Points indicate means and error bars indicate standard deviations across 100 replicate populations, and relative point size represents the proportion of populations surviving until the end of the simulation.

## Discussion

The results presented here illustrate that long-term bet-hedging strategies for choosing constant payoffs can outcompete short-term variance-prone strategies providing more variable payoffs, even though such variance-proneness offers higher arithmetic mean fitness. The importance of such bet-hedging effects depends upon the relative importance of additive versus multiplicative fitness accumulation (see Haaland et al. 2019). Our models allow individuals to accumulate fitness in the form of resources or reproductive success additively across *n* decision events within their lifetimes. However, the models also allow variation in the grain of the environment, which alters the correlations in fitness among individuals, and this in turn alters the extent to which long-term fitness of a genotype is determined additively versus multiplicatively across generations (Frank 2011; Starrfelt and Kokko 2012, Haaland and Botero *submitted*). Our simulation results broadly follow our intuition from Fig. 1 based upon existing theory, in that we observe interactive effects of the number of decision events (*n*) and the grain of the environment (*g*_*e*_) on the evolution of variance-averse versus variance-prone strategies.

In Model 1, where the only environmental parameter is the correlation in payoffs among individuals that choose the variable strategy, our simulated populations evolved a choice for the constant strategy when there were fewer decision events for accumulating resources prior to reproduction, and higher correlations in the variable payoffs between different individuals (Fig. 3). Our analytical calculations (Fig. 2) match our simulation results for *g*_*r*_ = 1 (i.e. when all individuals playing the variable strategy obtain the same reward), which shows that at *n* = 5 the two strategies are competitively equal in terms of long-term fitness. For *n*=1, this model captures the basic setup of well-known bet-hedging scenarios, such as that of annual plants in wet versus dry years (Seger and Brockmann 1987; Starrfelt and Kokko 2012; Crowley et al. 2016). Conservative bet-hedgers, coping moderately well with both wet and dry years, correspond to our individuals choosing the safe option, gaining a constant but moderate payoff no matter the state of the environment. Strategies specializing on one of the environments, essentially gambling that the environment will be the one that suits them best, correspond to our individuals choosing the variable option, gaining high enough payoffs when the environment is in ‘good’ condition (i.e. it matches their phenotype) that these cases outweigh the relative losses during ‘bad’ environmental conditions (i.e. when their phenotype is mismatched). In line with previous theory, when fitness accumulation is primarily multiplicative, bet-hedging dominates the outcome.

In Model 2, we introduced the mechanism that traditional variance-sensitivity literature assumes will favor variance-proneness: a sigmoid utility function relating individual energetic state to fitness and the predictions regarding variance sensitivity from the energy budget rule (Caraco 1981; Stephens 1981; Bednekoff 1996; Wright and Radford 2010; Kacelnik and El Mouden 2013). In this model, fitness correlations among individuals can come from two sources: correlations in environmental conditions determining energetic state (*g*_*e*_), and correlations in payoffs for individuals playing the variable strategy (*g*_*r*_). Here we found that variance-aversion is only favored when *n* is low (1 and arguably 2) and both *g*_*e*_ and *g*_*r*_ are high (0.75 or above). The relatively weaker selection for variance-aversion in this model compared to in Model 1 is not due to the arithmetic mean fitness benefits of variance-proneness being larger than in Model 1 (variance-proneness offers an increase in expected fitness of about 5-10 % for *x* < 0.5), but rather due to there being two uncorrelated sources of stochasticity affecting correlations in payoffs among individuals. For example, even if all individuals being variance-prone receive the same payoff (a scenario strongly favoring the ‘safe’ strategy in Model 1), variance-prone individuals may differ so much in energetic state that some individuals will still do well enough to ensure high genotype fitness despite the increase in fitness variation.

Interpreting our different modeling scenarios as representing the evolution of traits related to different activities, we can make simple inferences on the importance of bet-hedging in determining the evolution of variance-sensitivity as a viable strategy in nature. In particular, bet-hedging is unlikely to play a role in the evolution of variance-sensitive foraging preferences, such as in small-scale foraging patch or habitat choice, as these decisions for both animals and plants represent essentially an infinitely large number of (or even a continuous sequence of) decision events (Dener et al. 2016). Even if group members using the variance-prone strategy gain similar payoffs (i.e. *g*_*r*_ is high) in any one time step (e.g. one day), the additive accumulation of foraging payoffs over the large number of decision events (or days) that are experienced before the total resources are accumulated to determine reproductive success ensures that the strategy giving largest arithmetic mean payoffs should be favored evolutionarily. Thus, our modeling exercise does not reveal a direct flaw in the traditional predictions of variance-sensitivity as it applies to optimal foraging theory, and apparent failures of this paradigm to explain observed outcomes of foraging experiments (see Kacelnik and Bateson 1997; Kacelnik and El Mouden 2013) are likely to lie elsewhere, such as in the application of inappropriate statistical measures of variance when assessing outcomes (see Shafir 2000).

In contrast to the many decision events involved in foraging, certain important key decisions in the life of organisms are made only once or a few times, and can offer similar variance sensitive scenarios of choosing between ‘safe but low gain’ versus ‘high risk / high gain’ options. For example, decisions related to seasonal migration, including timing, choice of route, choice of destination, or even whether to migrate at all (Hebblewhite and Merrill 2009; Bauer et al. 2011; Grist et al. 2017; Reid et al. 2018), which are likely to include strong bet-hedging elements if there are correlations in the payoffs among related individuals that all choose the risky strategy. Therefore, ongoing rapid changes in the spatial scale of synchrony in resource availability or environmental variation as a consequence of anthropogenic climate change, habitat change or habitat destruction can all have dramatic effects on population viability (Shuter et al. 2011; Koenig and Liebhold 2016; Walter et al. 2017). The extent to which populations are able to adjust their strategy use in response to these changes is largely unknown, but it seems likely that such changes in fitness correlations among individuals are difficult to detect, so that previously advantageous variance-proneness may become an evolutionary trap (Robertson et al. 2013; Sih 2013).

Similarly, our model results suggest that bet-hedging may be a considerable selective force favoring variance-aversion in reproductive decisions and life-history strategies. For example, recent theoretical and comparative studies have suggested that cooperative breeding and cooperation in general might be adaptive because they offer lower variance in fitness returns in variable environments (despite the short-term reductions in expected fitness) as compared to solitary breeding and more selfish behavior (Rubenstein 2011; Koenig and Walters 2015; Shen et al. 2017; Kennedy et al. 2018). Our analyses broadly support these results in that bet hedging may be an important selective force when fitness correlations among individuals are high (i.e. coarse environmental grain, which emphasizes geometric mean fitness), but also demonstrate how additive fitness accumulation over the course of repeated variance-sensitive trials quickly shifts the balance to favor variance-proneness. Even with a coarse environmental grain, predictions for bet hedging through variance-averse reproductive decisions and life-history strategies will differ depending on certain (evolved) properties of the species, such as expected lifespans and degree of iteroparity (Hintze et al. 2015). Notably, a variance-prone reproductive strategy may still be favored in long-lived species where an individual can expect to breed many times throughout its life, given that the environmental variable determining whether this risky strategy is successful or unsuccessful is uncorrelated among breeding seasons (Bårdsen et al. 2008; Monteith et al. 2013). We therefore suggest that more progress can be made in understanding the evolution of cooperative breeding by not only studying the severity or magnitude of environmental fluctuations (Gonzalez et al. 2013; Cornwallis et al. 2017), but also how they affect the spatial and temporal scale of correlations in fitness payoffs.

Other life-history traits may respond similarly to the axes we outline here of spatiotemporal variation affecting arithmetic versus geometric mean fitness accumulation. For example, apparently sub-optimal clutch sizes in many bird species could represent a possible conservative bet-hedging strategy in response to year-to-year variation in spring onset and food availability (Boyce and Perrins 1987; Haaland et al. 2019), if those annual fluctuations are experienced by the entire population (i.e. a coarse environmental grain). This follows from larger clutches leading to higher reproductive success in good years and higher arithmetic mean fitness across years, but smaller clutches having much higher offspring survival in bad years, and thus a higher geometric mean fitness across years. Similar mechanisms are likely to be at play in the evolution of life span and/or body size in response to fluctuating density-dependent selection (Sæther et al. 2016; Wright et al. 2019), as well as the evolution of plasticity in reproductive attempts and effort. As the strength of fluctuating selection increases, and fitness correlations among individuals increase, larger costs related to informed plastic changes such as information gathering, learning and phenotypic adjustment are tolerated (Snell-Rood and Steck 2018; Wright et al. 2019). This dependence on among-individual correlations relates to differences among populations in the degree to which environmental stochasticity or demographic stochasticity drives population fluctuations. The relative importance of these can be estimated from field data (Lande et al. 2003), so this approach therefore identifies a starting point for examining the evolution of variance-sensitivity versus conservative bet-hedging in all types of life-history strategies.

In summary, our models support the growing understanding that bet-hedging strategies that reduce the variance in fitness can be favored in unpredictably varying environments despite reducing arithmetic mean fitness We demonstrate that variance-aversion can be an adaptive conservative bet-hedging strategy even when traditional variance-sensitivity theory predicts that variance-proneness should be favored. We also show that increasing the number of decision-making events over which payoffs accumulate will favor variance-proneness, and whilst this perhaps validates the approach taken in optimal foraging theory, it does have implications for the evolution of variance-sensitive life-history strategies. We also highlight the importance of environmental grain and the correlation of fitness pay-offs across individuals of the same genotype, which although common in bet-hedging theory is rarely considered in connection with variance sensitivity. We therefore hope that these ideas linking areas of theory in what have been quite disparate fields of study will improve to our understanding of potential evolutionary responses to environmental stochasticity and human-induced environmental change.

## Acknowledgements

We thank Steinar Engen and Gunnar A. Sveinsson for mathematical advice. TRH and IIR are supported by the Research Council of Norway on grant 240008 awarded to IIR on the Young Talented Researchers program, and this work was partly funded through its Centres of Excellence funding scheme, project number 223257, to Centre for Biodiversity Dynamics (CBD) at the Norwegian University of Science and Technology (NTNU).

